# Radiation exposure induced blood-brain barrier injury via mitochondria-mediated sterile inflammation

**DOI:** 10.1101/2025.02.03.636229

**Authors:** Peng Wang, Jiayue Liu, Min Zhang, Juan Yang, Peihan Lian, Xiu Cheng, Jianhua Qin

## Abstract

Radiation-induced brain injury (RIBI) is caused by exposure to high doses of ionizing radiation and characterized by severe cognitive dysfunction and brain necrosis. The widespread application of radiotherapy and rapid development of deep space exploration have substantially increased the risk of RIBI. However, the pathogenesis of RIBI is not fully understood, and no effective intervention is available. This work described a blood-brain barrier (BBB) microphysiological system (MPS), that allowed to explore the responses of BBB and distinct brain cells to radiation exposure. This BBB MPS can recapitulate the interface structure and function of BBB in brain microenvironment, including brain endothelial cells, astrocytes, pericytes and microglia co-cultured under flow condition. Following acute exposure to radiation of X-ray or γ-ray, characteristic RIBI-associated pathological responses were observed, including BBB compromise, DNA breaks, inhibited cell proliferation, cell hypertrophy and pro-inflammatory cytokine release. Among the distinctive types of cells, brain endothelial cells showed the highest radiosensitivity as compared to other cells in the MPS. Intriguingly, X-ray and γ-ray radiation consistently induced prominent sterile inflammation responses, especially type I interferon response, in the BBB MPS. These responses were mediated by radiation-induced mitochondrial DNA release and subsequent activation of cGAS-STING signaling pathway. Furthermore, we found abrocitinib (JAK1 inhibitor) and idebenone (mitochondrial protectant) can attenuate radiation-induced inflammation and ameliorate BBB injury. These findings revealed the involvement of mitochondria-mediated inflammation in RIBI pathogenesis, identifying mitochondria as a potential target for new radioprotective measures.

## Introduction

Radiation-induced brain injury (RIBI) refers to brain tissue injuries caused by exposure to high-dose ionizing radiation, which are very common in patients with head and neck tumors after radiotherapy ^1–4^. It can also occur in some extreme conditions, such as nuclear accidents or deep space exploration ^5,6^. Such injuries involve various pathological changes, including genomic damage, oxidative stress, decreased neurogenesis, glial cell activation, demyelination, neuroinflammation and cerebrovascular injury ^7–10^. However, the pathogenesis of RIBI is not fully understood, and no effective treatment is available.

Brain is one of the most vital human organs, in which blood-brain barrier (BBB) is a highly active barrier system to maintain the brain homeostasis. The BBB selectively regulates the transport of substances between blood circulation and brain tissue, and prevents pathogen invasion into the brain parenchyma ^11^. Studies have shown a close link between BBB dysfunction and the pathogenesis in some neurological diseases, such as Alzheimer’s disease and ischemic stroke ^12,13^. Besides, evidence increasingly indicates that brain vascular injury is a major pathological feature of RIBI ^14–18^. Studies have implicated the cerebral vasculature might be a primary radiation injury site, however, the mechanism behind these pathological changes remains elusive.

Conventionally, studies on RIBI have often relied on animal models, especially rodents, which have greatly advanced the research in radiation toxicology and radiation medicine ^14,19–22^. However, with the rising cost of animal experiments and the increasing concern for animal welfare and ethical issues, gradually reducing the use of animals is becoming the consensus of the academic community. In addition, due to differences in genetic background and physiological structure with human, animals often fail to predict the actual responses of human body to environmental stimulus or drugs. It’s needed to develop alternatives to conventional animal models that enables accurate assessment of the human body’s responses to radiation.

Microphysiological system (MPS), is a rapidly developing biotechnology that enables to emulate the functional units of human organs in a near physiological manner. It has been widely utilized in biomedical research, including disease modeling, drug testing, toxicological assessment, et al ^23,24^. Compared with the traditional 2D cell culture and animal models, MPS technology has showed unique advantages in recapitulating the brain physiological microenvironment, including the complex brain cells types, flow condition, ECM environment, etc. So far, various types of brain MPSs, such as BBB models or neurovascular unit models have been established, and are widely used in neuroscience research, including neurodegenerative diseases, ischemic stroke, brain infection and personalized medicine applications ^25–30^.

In this study, we developed a BBB MPS lined with brain endothelial cells, astrocytes, pericytes and microglia co-cultured under dynamic culture conditions to probe the effects of ionizing radiation. Exposure to X-ray or γ-ray radiation, it reveals the obvious injuries in the BBB MPS, including BBB leakage, DNA breaks, decreased cell density, cell-cycle arrest, cell hypertrophy and pro-inflammatory cytokine release. Radiation exposure also triggered substantial sterile inflammation, characterized by a remarkable type I interferon (IFN) response. This was also consistently detected in brain samples from RIBI patients ^31^. A mechanism study showed that the sterile inflammation was mediated by mitochondrial DNA release and subsequent cGAS-STING signaling pathway activation. Idebenone (mitochondrial protectant) or abrocitinib (type I IFN pathway inhibitor) pretreatment significantly ameliorated radiation-induced inflammation and BBB injury. We thus demonstrated the vital role of mitochondria-mediated inflammation in RIBI pathogenesis and identified the BBB MPS as a useful platform for developing novel RIBI therapeutics.

## Results

### Construction of a microengineered BBB MPS to probe the effects of radiation exposure

Radiation exposure is a risk factor that severely threaten the health of human brain, which happens in radiotherapy for head and neck tumors, nuclear accidents and deep space exploration (Fig. 1a). In this study, we designed a BBB MPS to probe the detrimental effects of radiation exposure on human BBB.

**Figure 1.**
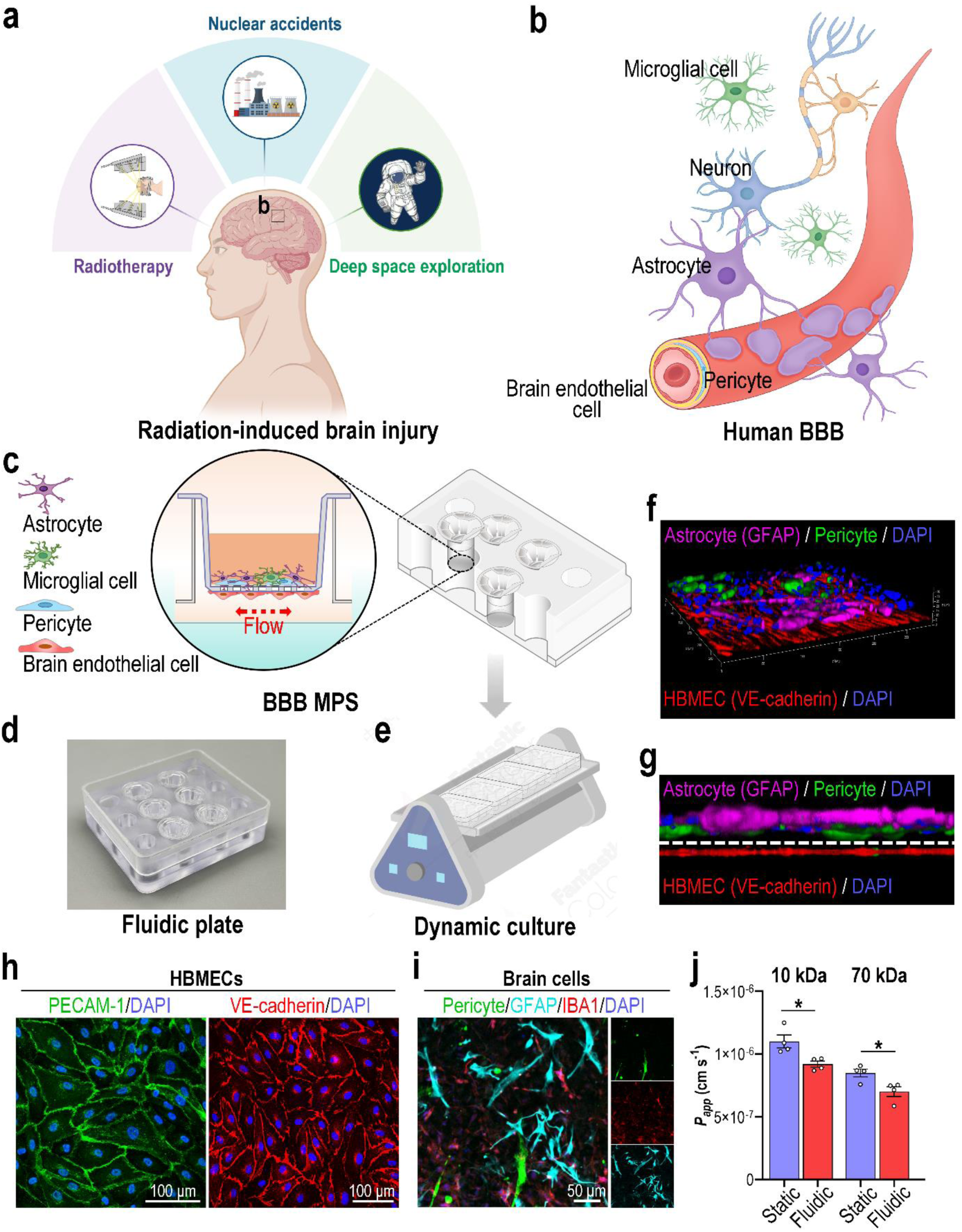
Establishment of a biomimetic BBB MPS on a fluidic culture plate. **a**, Schematic showing images of a human brain exposed to ionizing radiation. **b**, Schematic showing the anatomical structure of the human BBB. **c**, Schematic showing the establishment of a BBB MPS on a fluidic culture plate. **d**, Image of a real fluidic plate. **e**, Schematic showing the dynamic culture of BBB MPS on a shaker. **f**, 3D configuration image showing the BBB interface formed by HBMECs (VE-cadherin; red), pericytes (labeled by GFP) and astrocytes (GFAP; magenta) after co-culture under fluidic condition for 3 days. **g**, Side image showing the BBB interface formed by HBMECs (VE-cadherin; red), pericytes (green) and astrocytes (GFAP; magenta). **h**, Confocal fluorescent micrographs showing HBMECs in the BBB MPS via immunostaining for PECAM-1 (green) and VE-cadherin (red). **i**, Confocal fluorescent micrographs showing brain-side cells, namely pericytes (green), astrocytes (GFAP; cyan) and microglia (IBA1; red). **f-i**, Representative images from three independent samples. **j**, BBB MPS grown under static or fluidic culture conditions for 3 days and subjected to dextran (10, 70 kDa) permeability assays (*n* = 4). Data are presented as the mean ± SEM; unpaired Student’s t-test, *: *P* < 0.05.

In vivo, human BBB is mainly composed of brain endothelial cells, pericytes, astrocytes and basement membrane (Fig. 1b). Besides, microglia, the resident immune cells in central nervous system (CNS), play important roles in maintain brain homeostasis and processing of neurological diseases. In the model, human brain microvascular endothelial cells (HBMECs) were seeded on the lower sides of porous (1 μm) transwell insert membranes, and pericytes (hPSC-PCs), astrocytes and microglia (ratio = 1:5:1) were seeded on the upper sides (Fig. 1c, d). To simulate the fluid shear forces acting on endothelial cells, the transwell inserts were transferred to a fluidic plate and cultured under fluidic conditions with shaking (swing angle: 5°; swing frequency: 2 Hz) (Fig. 1e).

After 3 days, 3D confocal reconstitution imaging revealed a BBB-like barrier interface on the fluidic plate (Fig. 1f, g). Immunofluorescent micrographs of PECAM-1 and VE-cadherin staining showed an intact endothelium on the lower porous membrane side (Fig. 1h). Pericytes (GFP), astrocytes (GFAP; cyan) and microglia (IBA1; red) were co-cultured to mimic brain parenchymal tissue (Fig. 1i). A dextran assay (10 and 70 kDa) was performed to evaluate the model’s barrier function. Compared with the static culture condition, the fluidic culture condition significantly decreased the permeability index (Fig. 1j), indicating establishment of a functional BBB MPS on the fluidic plate.

### Radiation exposure induced obvious damage and inflammation responses in BBB MPS

We exposed BBB MPS to 16 Gy X-ray or γ-ray to probe the effects of ionizing radiation. 4 days later, a permeability assay (10 and 70 kDa dextran) revealed increased permeability of the exposed BBB MPS, indicating compromised BBB integrity (Fig. 2a-b). Consistent with the permeability assay, immunofluorescence analysis of VE-cadherin (Fig. 2c) showed a loss of junction continuity at cell boundaries and increased cell-free gaps (indicated by white arrows) in the brain endothelium of irradiated BBB MPS. Significant cell hypertrophy and decreased cell density were also detected in irradiated brain endothelial cells (Fig. 2d-e). Distinct responses were also observed among the three types of brain cells (Fig. 2f). Radiation exposure did not affect pericyte coverage (indicated by pericyte-labelled GFP in Supplementary Fig. 1) or astrocytic activity (indicated by GFAP in Supplementary Fig. 2). However, it induced obvious microglial activation, indicated by an increased IBA1 level (Supplementary Fig. 3). A significant decrease in cell density was also detected in the irradiated brain cells (Fig. 2g). To clarify whether this decrease was due to increased cell death, we assessed cell viability using TUNEL staining and found that radiation exposure did not induce significant cell death in BBB MPS (Supplementary Fig. 4).

**Figure 2.**
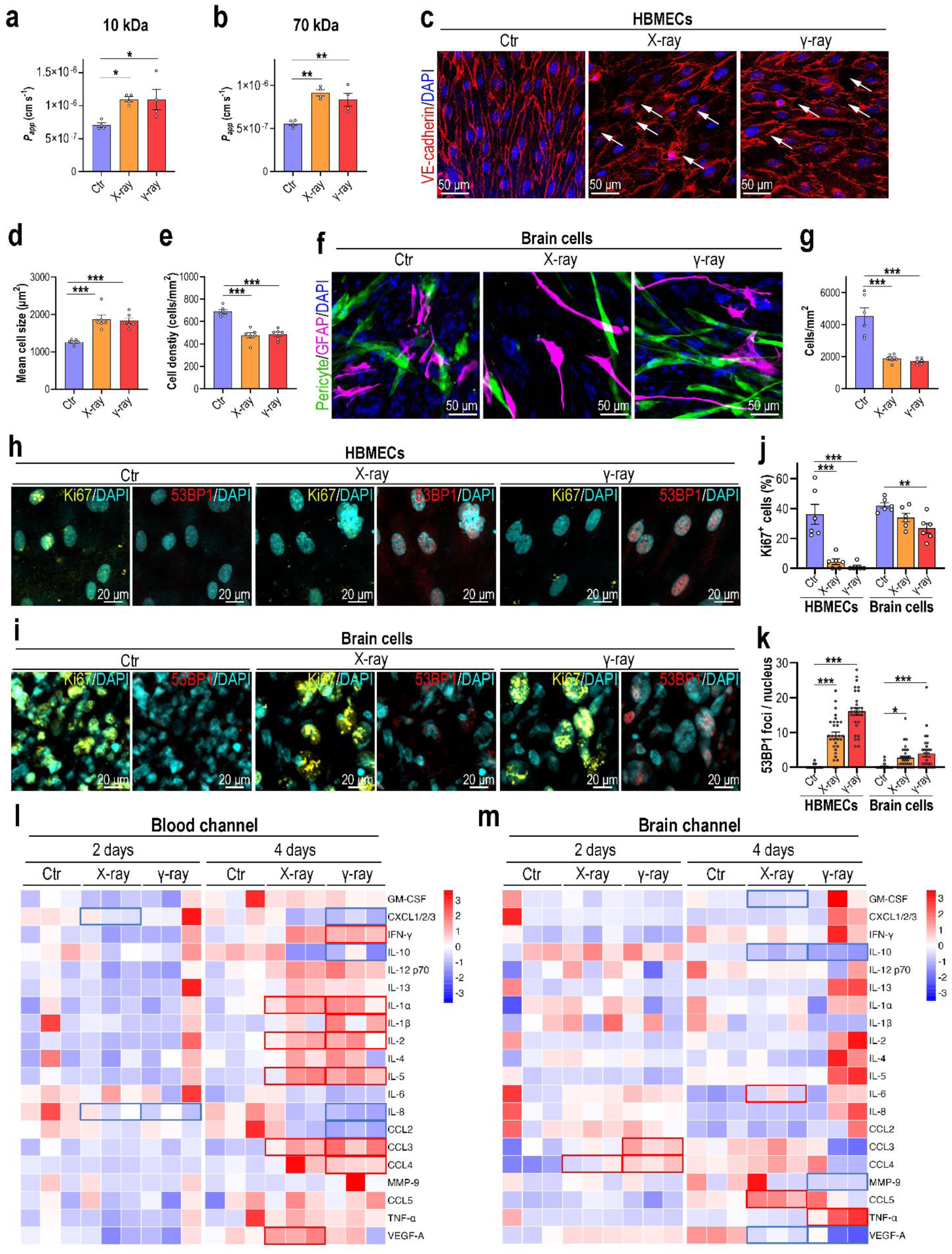
Exposure of the BBB MPS to radiation. **a**-**b**, Dextran (10, 70 kDa) assays performed to detect changes in BBB MPS permeability following 16 Gy X-ray or γ-ray radiation (*n* = 4). Data are presented as the mean ± SEM (unpaired Student’s t-test, *: *P* < 0.05). **c**, Confocal micrographs showing the HBMECs of BBB MPS 4 days after 16 Gy X-ray or γ-ray radiation (*n* = 3). **d**, Quantification of HBMEC size based on **c**. **e**, Quantification of HBMEC density based on **c**. **f**, Confocal micrographs showing pericytes (GFP) and astrocytes (GFAP; magenta) in BBB MPS 4 days after 16 Gy X-ray or γ-ray radiation (*n* = 3). **g**, Quantification of brain cell density based on **f**. **h**, Confocal micrographs showing HBMECs in BBB MPS detected by immunostaining for Ki67 (yellow) and 53BP1 (red) at 4 days post-radiation exposure (*n* = 3). **i**, Confocal micrographs showing brain cells in BBB MPS by immunostaining for Ki67 (yellow) and 53BP1 (red) at 4 days after radiation (*n* = 3). **j**, Quantification of percentages of Ki67^+^ cells in BBB MPS based on **h** and **i**. **k**, Quantification of 53BP1 foci per cell in BBB MPS based on **h** and **i**. **a**-**b**, **d**-**e**, **g**, **j**-**k**, Data are presented as the mean ± SEM (one-way analysis of variance followed by the Bonferroni post-hoc test, *: *P* < 0.05, **: *P* < 0.01, ***: *P* < 0.001). **l**, Cytokine array showing cytokine levels in the blood channels of BBB MPS at 4 days post-radiation exposure (*n* = 3). **m**, Cytokine array showing cytokine levels in the brain channels of BBB MPS at 4 days post-radiation exposure (*n* = 3). **l-m**, Downregulated cytokines with fold changes < 0.83 and *P* < 0.05 are indicated by blue boxes; upregulated cytokines with fold changes > 1.2 and *P* < 0.05 are indicated by red boxes. *P* values were calculated using a moderated t-test.

Radiation exposure might reduce cell density via cell proliferation. Ionizing radiation severely damages cells by generating DNA double-strand breaks (DSBs). We then assessed cell proliferation via immunostaining for Ki67 (a marker for cell proliferation) and DNA injury via immunostaining for 53BP1 nuclear foci (a marker of DSBs). Radiation exposure both significantly inhibited cell proliferation and induced DNA DSBs, and these effects were most severe in brain endothelial cells (Fig. 2h-k). Combined with the TUNEL assay results, ionizing radiation appeared to reduce cell density in BBB MPS by inducing DNA injury and inhibiting cell proliferation.

Inflammation is another characteristic of RIBI. We evaluated the inflammatory status of BBB MPS exposed to radiation using a multiple cytokine array. At day 4, the levels of several pro-inflammatory cytokines and growth factor, including IFN-γ, interleukin (IL)-1α, IL-1β, IL-2, IL-5, CCL3, CCL4 and VEGF-A, were significantly elevated in the blood channel following X-ray or γ-ray radiation (Fig. 2l). Lower levels were detected in the brain channels of irradiated BBB MPS (Fig. 2m). These results suggest that radiation exposure induced inflammation among the brain endothelial cells.

Collectively, the results indicate that either X-ray or γ-ray radiation of BBB MPS causes substantial damage, including disrupted BBB integrity, cell hypertrophy, decreased cell density, DNA damage, microglial activation and pro-inflammatory cytokine release. The BBB dysfunction we detected in the MPS was consistent with findings from the irradiated brain tissues in vivo ^8,9,18,32^, indicating that the BBB MPS could faithfully mimic the human brain’s pathological responses to radiation in vitro. Brain endothelial cells were more susceptible than other brain cells to ionizing radiation, indicating that cerebral blood vessels might be a major target in RIBI.

### Transcriptomic analysis revealed distinguished responses in brain endothelial cells and brain cells to radiation exposure

To better understand the underlying molecular signatures, we subjected BBB MPS at day 4 post-radiation exposure to transcriptomic analysis. In X-ray irradiated BBB MPS, 684 genes (359 upregulated, 325 downregulated) were dysregulated in brain cells, and 579 genes (158 upregulated, 421 downregulated) were dysregulated in brain endothelial cells (Fig. 3a). In γ-ray irradiated BBB MPS, 806 genes (400 upregulated, 406 downregulated) were dysregulated in brain cells, and 1846 genes (444 upregulated, 1402 downregulated) were dysregulated in brain endothelial cells (Fig. 3a). By comparing differentially expressed genes (DEGs) between irradiated brain endothelial and brain cells, we found only 64 and 145 DEGs shared in X-ray irradiated and γ-ray irradiated BBB MPS, respectively (Fig. 3b). That is, brain cells and brain endothelial cells exhibited distinct transcriptomic responses to either X-ray or γ-ray radiation. Then, we compared the transcriptomic responses of BBB cells to different types of ionizing radiation. Intriguingly, many genes were commonly modulated following X-ray and γ-ray radiation: 505 and 532 DEGs in the irradiated brain endothelial and brain cells, respectively (Fig. 2c). This finding suggests that X-ray and γ-ray radiation affect the human BBB similarly.

**Figure 3.**
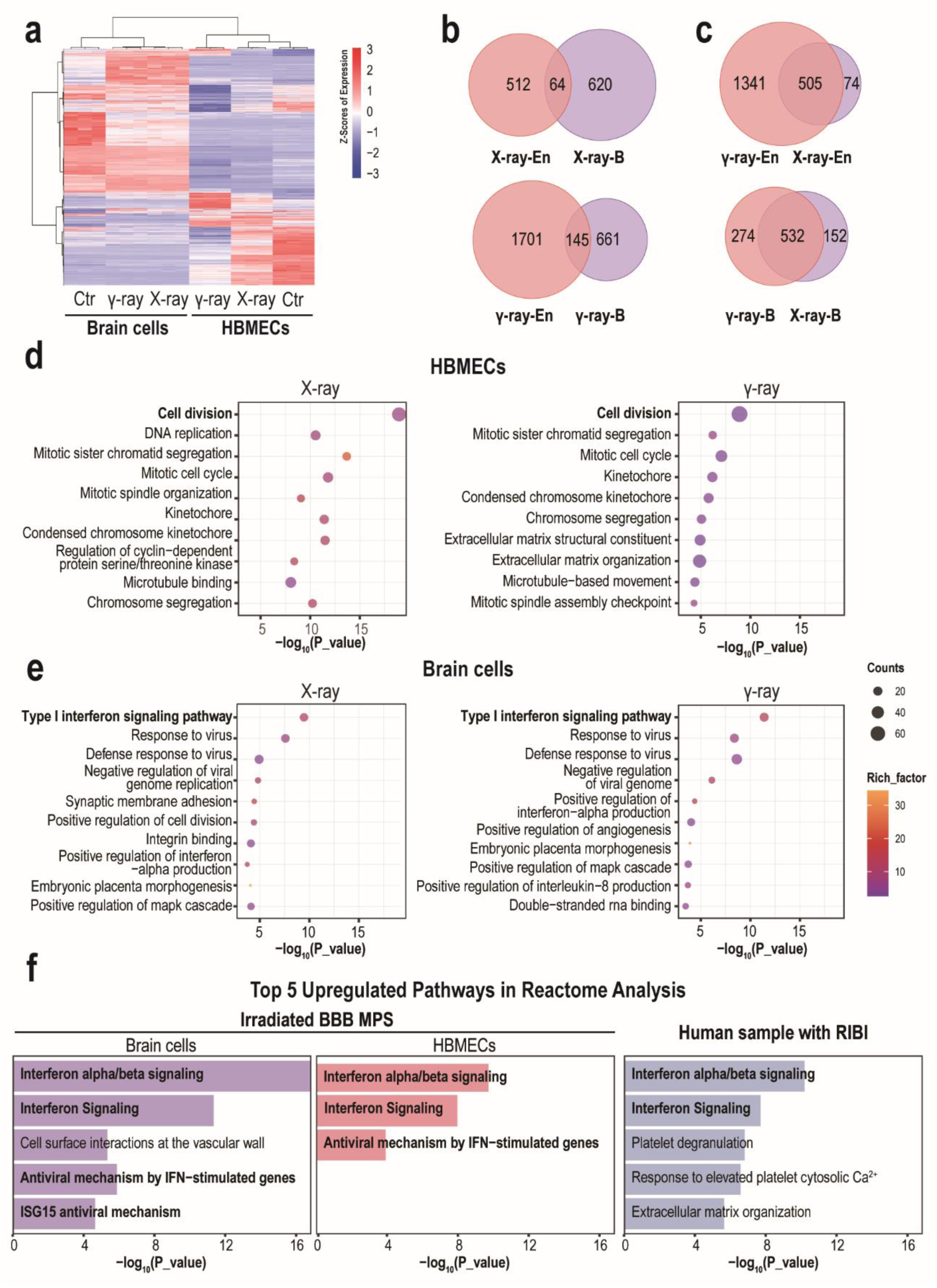
Transcriptomic analysis of cells on BBB MPS following radiation exposure. **a**, Heatmap showing differentially expressed genes (DEGs) in brain endothelial and brain cells at 4 days post-radiation exposure. Genes differentially expressed with fold changes of > 2.0 and *P* < 0.05 are defined as DEGs. *P* values were calculated using a two-sided, unpaired Student’s t-test with equal variance assumed (*n* = 3). **b**, Venn diagrams showing overlapping DEGs between HBMECs and brain cells following X-ray or γ-ray radiation. **c**, Venn diagrams showing overlapping DEGs between X-ray-and γ-ray-treated samples. **d**, GO enrichment analysis of DEGs in HBMECs following X-ray or γ-ray radiation. **e**, GO enrichment analysis of DEGs in brain cells following X-ray or γ-ray radiation. **f**. Reactome pathway enrichment analysis showing activated pathways in brain samples from patients receiving radiotherapy and irradiated BBB MPS. The RNA-seq data of brain samples from patients were obtained from a previous study ^31^.

Closed examination of DEGs in brain endothelial cells revealed that many genes related to vascular functions such as cell junctions, cell adhesion, angiogenesis and transport were significantly downregulated following radiation exposure, especially γ-ray irradiation (Supplementary Fig. 5). The brain endothelium plays a key role in BBB maintenance; ionizing radiation may cause a BBB break by damaging brain endothelial cells. We also analyzed the states of distinct brain cells using cell-specific markers. Gene expression analysis showed no increases in some classical astrocytic markers (e.g., GFAP, S100β, AQP4 and ALDH1L1; Supplementary Fig. 5), indicating no obvious astrocyte activation on the irradiated BBB MPS. However, the microglia-specific genes IBA1, CD45, CD68 and CD11b were significantly upregulated following radiation exposure (Supplementary Fig. 5), consistent with the increased IBA1 expression detected in irradiated BBB MPS by immunofluorescent analysis (Supplementary Fig. 3). Regarding pericytes, PDGFRβ was downregulated and NG2 was upregulated in irradiated BBB MPS (Supplementary Fig. 5).

Next, a detailed Gene Ontology (GO) enrichment analysis was performed to identify the biological processes underlying RIBI. In the irradiated brain endothelial cells, many DEGs were enriched in cell cycle-related processes (Fig. 3d) such as cell division, mitotic cell cycle, DNA replication and chromosome segregation. Gene set enrichment analysis (GSEA) revealed significantly inhibited cell division in irradiated brain endothelial cells (Supplementary Fig. 6), which explains the remarkable inhibition of cell proliferation in the irradiated brain endothelial cells (Fig. 2h, j). GO enrichment analysis of irradiated brain cells was also performed, revealing enrichment of DEGs in some inflammation-related processes, especially anti-viral responses such as the type I IFN signaling pathway, positive regulation of IFN-α production, defense responses to virus and negative regulation of viral genome replication (Fig. 3e).

Taken together, transcriptomic analyses revealed that X-ray and γ-ray radiation similarly induce BBB injury, whereas responses to ionizing radiation are cell type-dependent.

### Type I IFN signaling pathway was activated in irradiated BBB MPS

As the transcriptomic analysis revealed type I IFN signaling pathway modulation in irradiated BBB MPS, we performed GSEA to analyze the potential correlation between this pathway and radiation exposure. We observed significant upregulation of the type I IFN pathway in both brain endothelial and brain cells from irradiated BBB MPS (Supplementary Fig. 7). We also performed Reactome pathway enrichment analysis to compare activated signaling pathways between patients’ brain samples and BBB MPS following radiation exposure. RNA-seq data of patients’ brain samples were obtained in a previous study ^31^. Intriguingly, similarly activated signaling pathways were detected in brain samples from patients receiving radiotherapy and irradiated BBB MPS, as reflected by enriched IFNα/β and IFN signaling (Fig. 3f). That is, radiation exposure activated type I IFN responses in human brains, verifying the reliability of the BBB MPS for modeling RIBI physiopathology in vitro. We further analyzed the expression of type I IFNs and their receptors by qPCR and found that radiation exposure upregulated IFNE in brain endothelial cells and IFN receptors (IFNAR1, IFNAR2) in brain cells (Fig. 4a). We also observed significant upregulation of the type I INF pathway genes STAT1, MX1, ISG15, IRF7 and IFIT3 in both brain endothelial and brain cells from irradiated BBB MPS (Fig. 4b).

**Figure 4.**
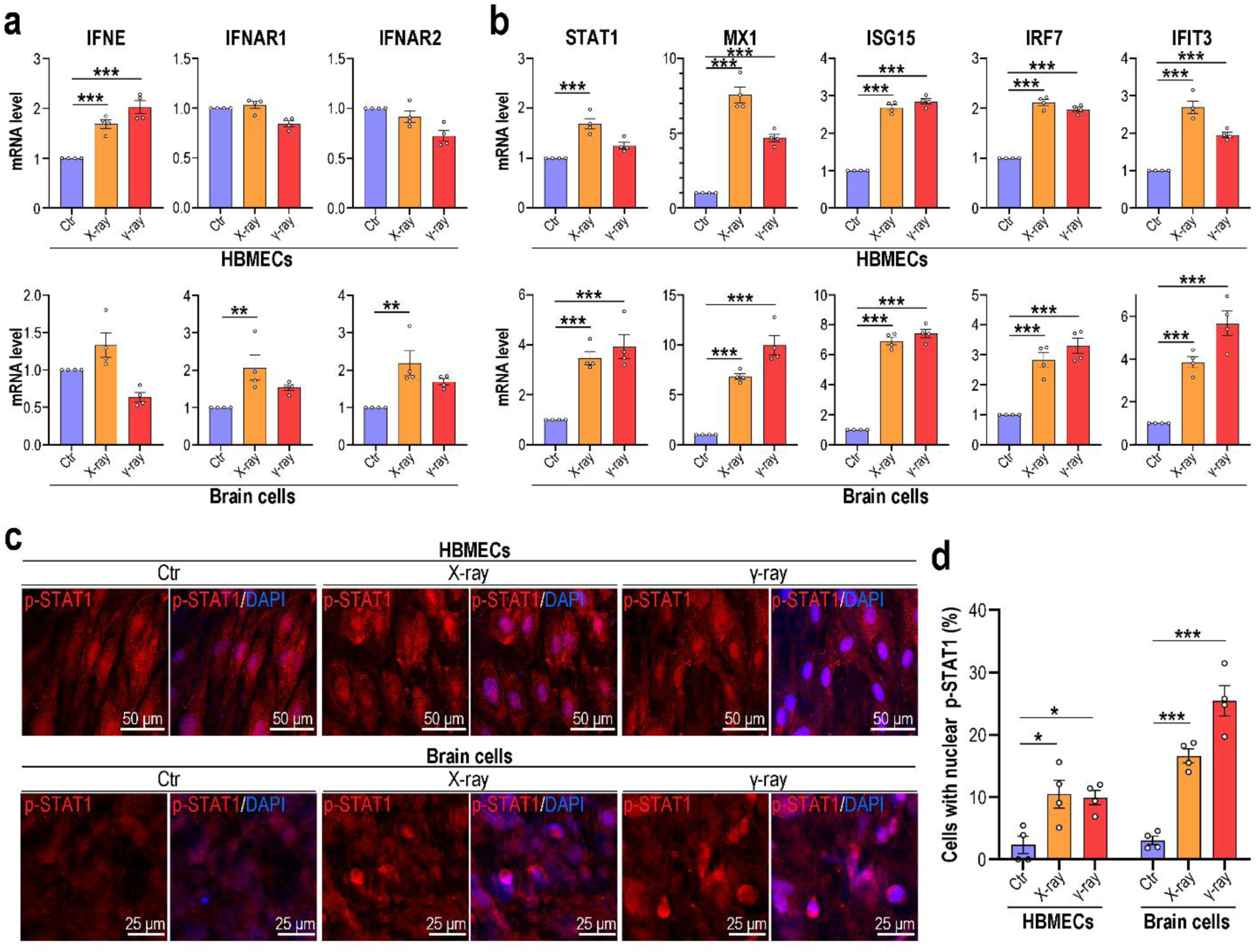
Radiation exposure induced JAK/STAT1 pathway activation in BBB MPS. **a**, Levels of IFNE and IFNAR (IFNAR1, IFNAR2). mRNA in HBMECs and brain cells detected by qPCR 4 days post-radiation exposure (*n* = 4). **b**, mRNA levels of genes associated with the type I IFN signaling pathway in HBMECs and brain cells detected by qPCR 4 days post-radiation exposure (*n* = 4). **a**-**b**, GAPDH was utilized as an internal control. The mRNA levels of target genes in control group are normalized as 1. **c**, Confocal micrographs of cells on BBB MPS immunostained for p-STAT1 at 4 days post-radiation exposure (*n* = 4). **d**, Quantification of cells with nuclear p-STAT1 localization based on **c**. **a**-**b**, **d**, Data are presented as the mean ± SEM (one-way analysis of variance followed by the Bonferroni post-hoc test; **: *P* < 0.01, ***: *P* < 0.001).

Type I IFNs are a vital class of cytokines that coordinate innate immune responses to infection, tumors and inflammation ^33^. Canonically, these cytokines activate the JAK/STAT signaling pathway, initiating the transcription of IFN-stimulated genes (ISGs) ^33–35^. We examined JAK/STAT signaling by detecting STAT1 (Ser727) phosphorylation, which enables the translocation of STAT1 from the cytoplasm to the nucleus and increases transcriptional activity ^34,35^. Immunofluorescence analysis showed a significant increase in p-STAT1 nuclear localization in both brain endothelial and brain cells from irradiated BBB MPS, indicating that ionizing radiation activates type I IFN signaling in the human brain.

### Radiation exposure elicited sterile inflammation via the mitochondrial DNA-mediated cGAS-STING pathway

Mitochondrial dysfunction and subsequent mitochondrial DNA (mtDNA) release can trigger innate immune responses involving the production of type I IFNs and other proinflammatory cytokines ^36,37^. We further investigated the modulatory role of mitochondria in the innate immune responses underlying RIBI. Using tetramethylrhodamine methyl ester (TMRM), a membrane potential-dependent mitochondrial fluorescent dye, and fluorescent micrography, we detected a remarkable decrease in mitochondrial membrane potential in brain endothelial cells post-radiation exposure but no obvious effects in brain cells (Fig. 5a-b). Together with the cytokine analysis, TMRM staining further indicated that radiation exposure affects brain endothelial cells more strongly than brain cells. Next, we used transmission electron microscopy (TEM) to examine the mitochondrial ultrastructure in brain endothelial cells from BBB MPS, revealing increases in over-fragmented mitochondria and mitochondria with vacuoles (yellow and red arrows, respectively, Fig. 5c) in irradiated brain endothelial cells.

**Figure 5.**
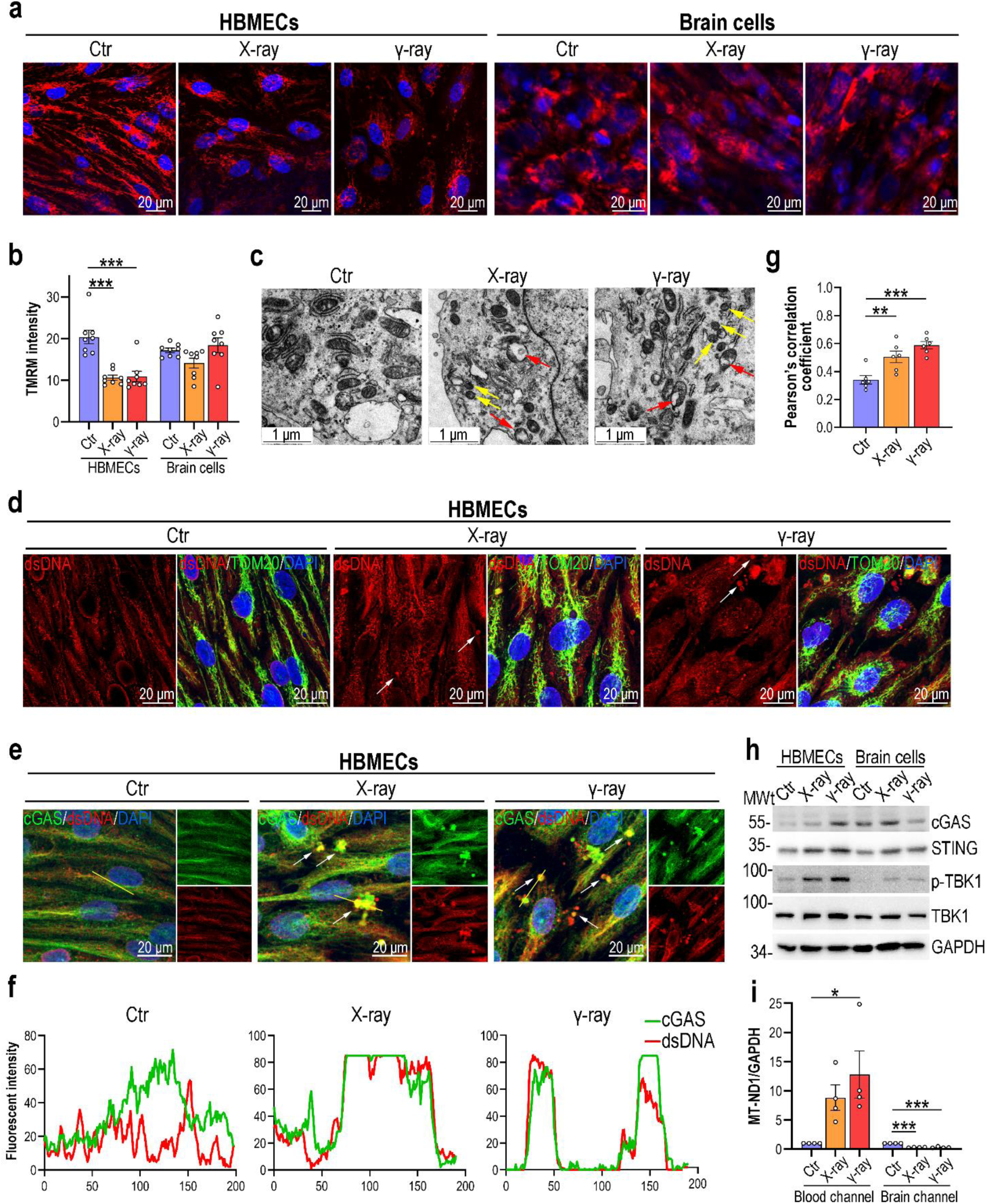
Radiation exposure induced mtDNA release and subsequent cGAS-STING pathway activation in the BBB MPS. **a**, Confocal micrographs of HBMECs and brain cells stained for TMRM at 4 days post-radiation exposure (*n* = 4). **b**, Quantification of TMRM fluorescence intensity in HBMECs and brain cells based on **a**. Two fields were quantified per sample. Data are presented as the mean ± SEM (one-way analysis of variance followed by the Bonferroni post-hoc test, ***: *P* < 0.001). **c**, TEM images showing the HBMEC ultrastructure at 4 days post-radiation exposure (*n* = 3). Mitochondria with swollen cristae and fragmented mitochondria are indicated by red and yellow arrows, respectively. **d**, Confocal micrographs of HBMECs immunostained for dsDNA (red) and TOM20 (green) at 4 days post-radiation exposure (*n* = 3). Cytoplasmic mtDNA aggregates are indicated by white arrows. **e**, Confocal micrographs of HBMECs immunostained for dsDNA (red) and cGAS (green) at 4 days post-radiation exposure (*n* = 3). dsDNA aggregates and cGAS colocalization signals are indicated by white arrows. **f**, Line-scan analysis of dsDNA (red) and cGAS (green) along the yellow line in **e**. **g**, Quantification of dsDNA (red) and cGAS (green) colocalization using Pearson’s correlation coefficients (*n* = 3). Two fields were quantified per sample. Data are presented as the mean ± SEM (one-way analysis of variance followed by the Bonferroni post-hoc test, *: *P* < 0.05, ***: *P* < 0.001). **h**, Western blot analysis showing the levels of cGAS, STING, p-TBK1 and TBK1 in HBMECs and brain cells following radiation exposure (*n* = 3). **i**, qPCR results showing the relative mtDNA copy numbers in medium from the blood and brain channels at 4 days post-radiation exposure (*n* = 4). GAPDH was utilized as an internal control. The relative copies of mtDNA in the control group are normalized as 1. Data are presented as the mean ± SEM (one-way analysis of variance followed by the Bonferroni post-hoc test, *: *P* < 0.05, ***: *P* < 0.001).

During mitochondrial injury, mtDNA can be released into the cytosol. We detected the sub-cellular localization of mtDNA using confocal microscopy. Co-immunostaining for TOM20 (mitochondrial marker) and DNA (using a dsDNA antibody) revealed an increase in DNA nucleoids in the cytosol of irradiated brain endothelial cells (Fig. 5d; Supplementary Fig. 8). The cGAS-STING signaling pathway can recognize released mtDNA as a foreign pathogen and initiate downstream immune responses ^36^. However, it is unclear whether RIBI triggers this response. We then detected cGAS-STING pathway activation in irradiated brain endothelial cells. First, we analyzed the co-localization of mtDNA and cGAS via co-immunostaining and found a significant increase in cytosolic co-localization in irradiated brain endothelial cells (Fig. 5e-g). We also used Western blotting to examine the expression of cGAS-STING pathway-associated proteins and revealed significant upregulation of cCAS, STING and TBK1 expression and TBK1 phosphorylation in irradiated brain endothelial cells (Fig. 5h). To test whether an increase in cytosolic mtDNA led to the increased extracellular release of mtDNA, we determined the level of cell-free mtDNA in BBB MPS medium using qPCR. The cell-free mtDNA copy number was significantly elevated in the blood channels of irradiated BBB MPS (Fig. 5i), indicating that the cell-free mtDNA level might be a diagnostic biomarker of RIBI.

To further confirm the above findings on mitochondrial dysfunction and cCAS-STING pathway activation in lesioned brain tissues from patients with RIBI, we re-analyzed previously published single-cell RNA sequencing (scRNA-seq) data ^19^. In brain endothelial cells, we observed many downregulated genes associated with mitochondria-related biological processes (Supplementary Fig. 9a), such as the respiratory electron transport chain, mitochondrial respirasome, mitochondrial inner membrane, mitochondrial protein-containing complex and ATP biosynthesis. Violin plots showed that many mtDNA-encoded genes, including MT-ND1, MT-NF2, MT-ND5, MT-ND4, MT-ND4L, MT-ATP6, MT-CO1, MT-CO2, MT-CO3 and MT-CYB, were significantly downregulated (Supplementary Fig. 9b) and TBK1 was upregulated in the brain endothelial cells of patients with RIBI (Supplementary Fig. 9c).

Collectively, these findings indicate that in brain endothelial cells, radiation exposure induces mitochondrial dysfunction and subsequent innate immune responses via the cGAS-STING signaling pathway.

### Abrocitinib and idebenone showed radioprotective efficacy for radiation-induced BBB injury

We next tested whether drugs targeting mitochondrial dysfunction or inflammation could modify radiation-induced BBB injury. We chose six components: hydrocortisone (anti-inflammatory), crisaborole (anti-inflammatory), abrocitinib (JAK1 inhibitor), coenzyme Q10 (antioxidant), BAI1 (BAX inhibitor) and idebenone (mitochondrial protectant). Four hours before radiation exposure, BBB MPSs were pre-treated with the indicated drugs; control and irradiated BBB MPSs were examined 4 days later (Fig. 6a). A BBB permeability assay (10 kDa-dextran) showed that both abrocitinib and idebenone significantly ameliorated radiation exposure-induced BBB compromise (Fig. 6b). Using immunofluorescence to determine the cellular state, we found that, consistent with the BBB permeability assay, abrocitinib or idebenone partially prevented endothelial injury in the irradiated BBB MPS (Fig. 6c), as indicated by decreased intercellular spaces between adjacent cells (Fig. 6d). However, brain endothelial cell hypertrophy was not significantly restored following abrocitinib or idebenone treatment (Fig. 6e). Microglia were also examined via immunostaining for IBA1: both abrocitinib and idebenone partially (but not significantly) restored IBA1 expression in the irradiated BBB MPS (Fig. 6f-g).

**Figure 6.**
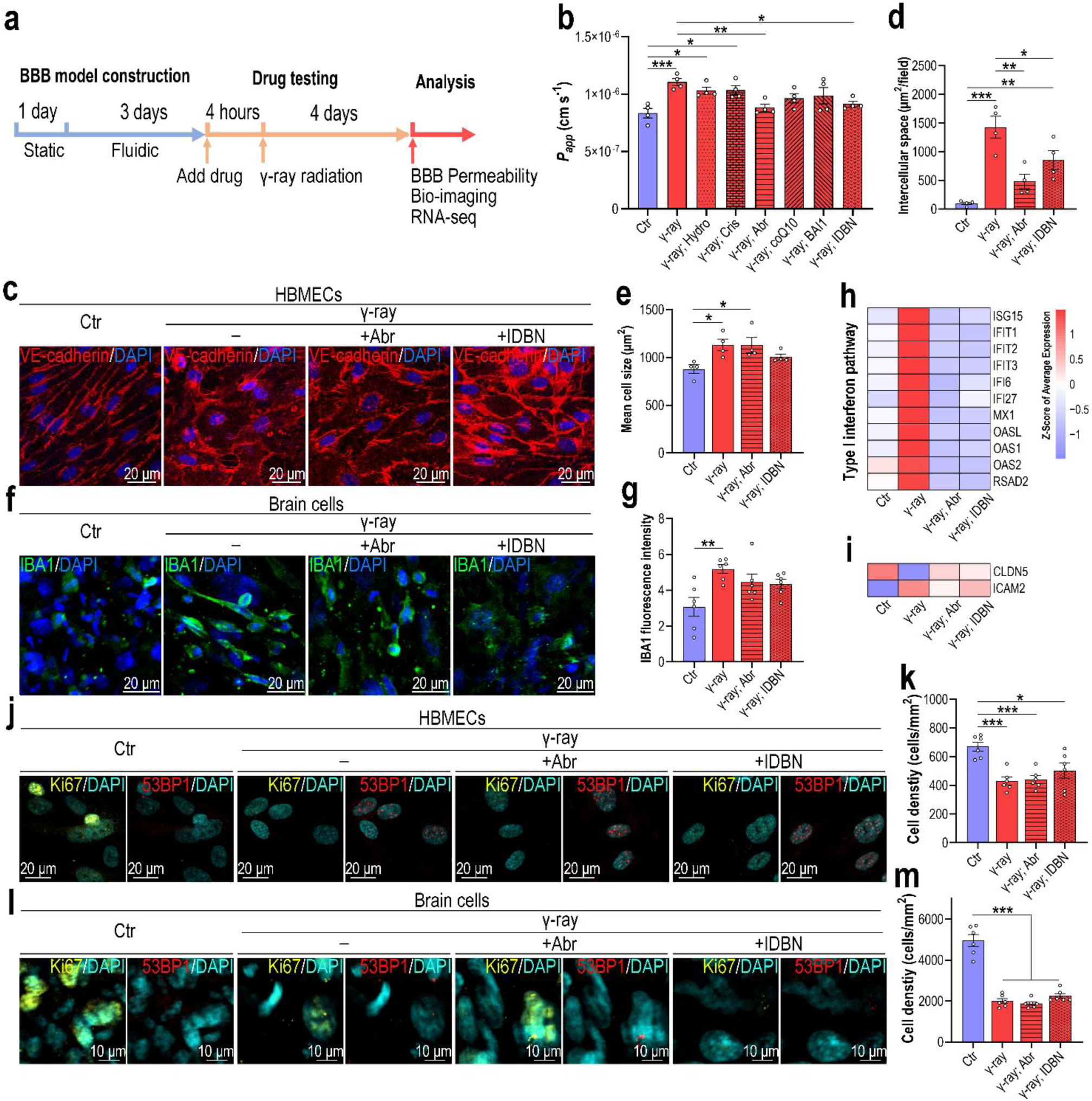
Drug testing in irradiated BBB MPS. **a**, Schematic showing the experimental drug testing procedure involving irradiated BBB MPS. **b**, A 10 kDa-dextran assay showing the permeability of irradiated BBB MPS following drug treatment (*n* = 4). Hydro: hydrocortisone; Cris: crisaborole; Abr: abrocitinib; coQ10: coenzyme Q10; BAI1: BAI1; IDBN: idebenone. **c**, Confocal micrographs of HBMECs from irradiated BBB MPS following abrocitinib or idebenone treatment (*n* = 4). **d**, Quantification of the intercellular space per field based on **c** (*n* = 4). e, Quantification of the mean HBMEC size based on **c**. **e**, Quantification of HBMEC density based on **c** (n = 4). **f**, Confocal micrographs of microglia from irradiated BBB MPS following abrocitinib or idebenone treatment (*n* = 4). **g**, Quantification of IBA1 immunofluorescence intensity in microglia based on **f** (*n* = 4). **h**, Heat map showing type I IFN pathway-related genes in irradiated HBMECs following abrocitinib or idebenone treatment (*n* =3). **i**, Heat map showing expression of CLDN5 and ICAM2 in irradiated HBMECs following abrocitinib or idebenone treatment (*n* =3). **j**, Confocal micrographs of irradiated HBMECs immunostained for Ki67 (yellow) and 53BP1 (red) following abrocitinib or idebenone treatment (*n* = 3). **k**, Quantification of HBMEC density based on **j** (*n* = 3). **l**, Confocal micrographs of irradiated brain cells immunostained for Ki67 (yellow) and 53BP1 (red) following abrocitinib or idebenone treatment (*n* = 3). **m**, Quantification of brain cell density based on **l** (*n* = 3). **b**, **d**-**e**, **g**, **k**, **m**, Data are presented as the mean ± SEM (one-way analysis of variance followed by the Bonferroni post-hoc test, *: *P* < 0.05, **: *P* < 0.01, ***: *P* < 0.001).

To identify the molecular basis of RIBI treatment via abrocitinib or idebenone, we subjected brain endothelial cells from BBB MPS to RNA-seq analysis. First, we examined the expression of genes related to innate immune responses. Consistent with previous findings (Supplementary Fig. 7), radiation exposure dramatically upregulated type I IFN signaling pathway-related genes in brain endothelial cells, while abrocitinib or idebenone treatment significantly restored gene expression to normal levels (Fig. 6h). Several upregulated leukocyte migration-related genes were also restored to normal levels following idebenone treatment (Supplementary Fig. 10). Intriguingly, the radiation-induced decrease in CLDN5 gene expression was also restored following abrocitinib or idebenone treatment (Fig. 6i).

As radiation exposure caused severe DNA damage and cell-cycle arrest in BBB MPS, we analyzed whether abrocitinib or idebenone could restore cell division at the transcriptomic level. However, neither drug effectively restored the expression of cell division-related genes (Supplementary Fig. 11). Immunostaining (Fig 6j, l) verified that neither abrocitinib nor idebenone could reverse DNA damage and cell cycle arrest in the irradiated BBB MPS, as revealed by the quantification of Ki67^+^ cells (Supplementary Fig. 12a, c), DSBs (Supplementary Fig. 12b, d), and cell density (Fig. 6k, m).

Collectively, these findings indicate that idebenone or abrocitinib can significantly mitigate radiation-induced brain endothelial injury and BBB dysfunction by repairing damaged mitochondria or blocking subsequent type I IFN pathway activation.

## Discussion

RIBI is a common, fatal complication of cranial radiotherapy, and no effective interventions are available. Here, we established a biomimetic human BBB MPS to recapitulate the basic structure and functional characteristics of BBB in vitro. The BBB MPS was exposed to X-ray and γ-ray irradiation and, intriguingly, exhibited similar responses to the two types of radiation, including increased BBB permeability, endothelial injury, decreased cell density, DNA breaks, cell-cycle arrest and pro-inflammatory cytokine release. RNA-seq analysis verified these findings, revealing a significant overlap in DEGs between X-ray and γ-ray irradiated cells, suggesting that BBB injuries following different irradiation types might share a molecular basis. From the perspective of drug development, this finding is favorable as it suggests that drugs may be effective against brain injuries caused by different types of radiation.

The human BBB comprises various distinctive cells, including brain endothelial cells, pericytes and astrocytes. These cell types play distinctive physiological roles to maintain normal BBB function. Following radiation exposure, different cell types exhibited distinctive susceptibilities: of the four types of cells in the BBB MPS, brain endothelial cells incurred the most damage. Following radiation exposure, more DNA breaks were detected in brain endothelial cells than other brain cells, and almost no proliferating cells were present in irradiated brain endothelium. This finding was verified by RNA-seq analysis, which revealed the dysregulation of many cell proliferation-related genes in irradiated brain endothelial cells. Pro-inflammatory cytokine release was mainly detected in the blood channel. As brain endothelium is the core component of BBB, we speculated that brain endothelial cell damage is closely associated with radiation-induced BBB comprise. The high sensitivity of endothelial cells to radiation exposure was previously reported in studies revealing severe endothelial injury and decreased brain capillary density in patients with RIBI and irradiated animal models ^17,21^. In one study, transplantation of endothelial progenitor cells reversed vascular injury and rescued cognitive function in irradiated mice, indicating the potential benefits of BBB-targeted therapy for RIBI ^38^.

We also detected prominent type I IFN pathway activation in the irradiated BBB MPS. This finding was unexpected, as type I IFN responses are generally thought to comprise an important defense mechanism against invading pathogens, especially viruses ^33,39,40^. Radiation exposure significantly triggered type I IFN responses in both brain endothelial and brain cells, including the upregulation of type I IFN and receptor genes and activation of the downstream JAK/STAT signaling pathway. This finding was validated in vivo via transcriptomic analysis of brain samples from glioblastoma patients treated with radiotherapy (Fig. 3f). As a class of cytokines with major anti-infection functions, type I IFNs must be strictly regulated; otherwise, they will induce serious abnormal immune responses. For example, many immune-mediated inflammatory diseases, such as systemic lupus erythematosus, systemic sclerosis, inflammatory myositis and Sjögren’s syndrome, are closely related to excessive type I IFN production ^41,42^. We speculate that radiation exposure triggers excessive inflammatory responses, especially type I IFN, which contribute to BBB injury.

Mitochondrial dysfunction can trigger mtDNA release into the cytoplasm; cytoplastic DNA sensors may mis-recognize this mtDNA as an invading foreign genome and initiate downstream innate immune responses ^36,37,43^. We then explored the potential relationship between type I IFN pathway activation and mitochondrial dysfunction. As expected, in irradiated brain endothelial cells, we detected obvious mtDNA release and subsequent cGAS-STING pathway activation, which is known to trigger innate immune responses involving type I IFN and other pro-inflammatory cytokine production ^36,44^. Consistent with mitochondrial dysfunction in the irradiated BBB MPS, scRNA-seq analysis of RIBI samples revealed the significantly dysregulated expression of many mitochondrial respiratory chain-associated genes and mtDNA-encoded genes in brain endothelial cells (Supplementary Fig. 9), indicating that brain endothelial injury in RIBI is dominated by mitochondrial dysfunction. Several studies have also reported radiation-induced mitochondrial dysfunction in cells ^45,46^. Our study reveals for the first time that radiation-induced mitochondrial dysfunction can elicit innate immune responses in the BBB by activating the cGAS-STING pathway.

Following RIBI mechanism study, we screened a few potential radioprotective drugs with anti-inflammatory or mitochondrial protective functions. Intriguingly, two drugs, abrocitinib and idebenone, had significant therapeutic effects on radiation-induced BBB injury, as evidenced by the partial recovery of intercellular junctions in the brain endothelium and decreased BBB permeability and inflammatory responses. Abrocitinib is an orally active selective JAK1 inhibitor that effectively treats some autoimmune diseases, including atopic dermatitis and mucous membrane pemphigoid ^47,48^. In this study, abrocitinib effectively attenuated radiation-induced BBB injury, indicating the vital role of type I IFN and the downstream JAK/STAT signaling pathway in disease pathogenesis. Idebenone is a well-appreciated mitochondrial protectant that protects against some neurological disorders in vivo, including ischemic stroke and Parkinson’s disease ^49,50^. Idebenone, a mitochondrial protective agent, significantly reduced the expression of inflammation-related genes, especially those related to type I IFN responses. This finding further demonstrates that mitochondrial damage is an early pathogenic factor triggered in the BBB by radiation exposure that initiates innate immune responses and eventually leads to BBB dysfunction.

This study still has some limitations. First, the parenchyma is a 3D tissue in the human brain; however, brain cells (pericytes, astrocytes and microglia) were cultured under 2D conditions in the BBB MPS, which may have affected their states. A previous study reported that 2D culture reduces the number of astrocyte processes and affects the polarized distribution of AQP4 in end-feet ^51^. Second, although they are not involved in BBB formation, neurons are the most vital cell type in the CNS and the functional basis of neural activity. In future work, an evaluation of how radiation-induced BBB dysfunction affects neuronal function and viability by incorporating neurons into the BBB MPS would be meaningful.

In summary, this study represents a new attempt to probe the effects of radiation on the human BBB model at tissue level. The unique advantage of this model is its ability to mimic the physiological structure and function of the BBB in vitro and capture pathological responses of different cells to radiation stimuli virtually. We discovered that excessive type I IFN response activation contributed to vascular injury following radiation exposure, which was mediated by mitochondrial dysfunction and subsequent cGAS-STING pathway activation. Importantly, we identified a mitochondrial protectant (idebenone) and JAK1 inhibitor (abrocitinib) that effectively ameliorated radiation-induced inflammation and BBB injury. Collectively, we uncovered an unknown mitochondria-mediated sterile inflammation involved in RIBI pathogenesis, which indicated mitochondria might be utilized as the therapeutic target for this devastating disease.

## Materials and methods

### Cell culture

HBMECs were purchased from ScienCell Corporation (Cat #: 1000) and cultured in endothelial cell medium (ECM; ScienCell; Cat #: 1001) supplemented with 5% FBS. HA cells were purchased from ScienCell Corporation (Cat #: 1800) and cultured in AM medium (ScienCell; Cat #: 1801). HMC3 cells were purchased from Procell Corporation (Cat #: CL-0620) and maintained in MEM medium containing NEAA (Procell; Cat #: MP150410) supplemented with 10% FBS and 1% P/S. hPSC-derived pericyte-like cells (hPSC-PCs) was a kind gift from Prof. Andy Peng Xiang’s lab (Sun Yat-Sen University) and were cultured in a PM medium (ScienCell; Cat #: 1201) supplemented with 50 ng/mL PDGF-BB (Peprotech; Cat #: 100-14B). hPSC-PCs were transfected with a lentivirus-GFP vector for 48 hours, and GFP^+^ hPSC-PCs were selected by 10 μg/ml puromycin dihydrochloride (Beyotime; Cat #: ST551). All cells were cultured at 37 °C in a humidified atmosphere of 5% CO_2_.

### Cell culture on the fluidic plate

Before seeding cells, transwell inserts (1 μm pore diameter; 2 × 10^6^ pores/cm^-2^; Greiner; Cat #: 662610) were pre-coated with 50 μg/mL fibronectin at 37 °C overnight. To construct the human alveolus chip, 100 μl of HBMEC cell suspension (200 cells/μl) was initially added on the bottom side of the transwell insert and allowed cells to attach to the membrane surface. Two hours later, transwell inserts were transferred to the 24-well plate, and 1 ml ECM medium was added in to the lower well. Then, 200 μl of brain cell mixture suspension (HA cells: 200 cells/μl; hPSC-PCs: 40 cells/μl; HMC3 cells: 40 cells/μl) was added into the upper well of the transwell insert. The brain medium was a mix of AM and PM medium (AM: PM = 1: 1), and the medium was changed every two days. After static culture for 1 day, transwell inserts were transferred to the high-throughput fluidic plate, and cultured under fluidic condition (tilting angle = 5°; rocking frequency = 2 Hz) at 37℃ for 3 days to form a confluent BBB interface.

### Radiation exposure

After fluidic culture for 3 days, fluidic plate was exposed to X-ray or γ-ray radiation at a dose of 16 Gy. The control samples (0 Gy) were removed from the 37℃ incubator and placed at room temperature inside the biosafety cabinet. Following exposure to radiation, all plates were transferred back to 37℃ incubator, and continued to culture under fluidic condition for another 4 days. The medium was changed every two days.

### Immunofluorescent imaging

Cells on the transwell chambers were washed twice with PBS and fixed with 4% PFA at 4 °C overnight. The fixed cells were permeabilized and blocked with 0.2% Triton X-100 in PBS (PBST) buffer containing 5% normal goat serum for 30 min at room temperature. Antibodies were diluted with PBST buffer and added into the upper compartment and lower compartment, respectively. Cells were stained with the primary antibodies at 4 °C overnight and corresponding secondary antibodies at room temperature for 2 h. Finally, the cells were counterstained with DAPI, and the transwell chambers were mounted on the coverslips. Images were acquired using a Leica STELLARIS 5 confocal fluorescent microscope. The Pearson’s *R* value was determined for immunofluorescent images using the ImageJ Colocalization Finder plugin.

### Multiplex assay for cytokine detection

Culture supernatant was harvested separately from the upper and lower channels of the chips. 20 cytokines in the culture supernatants were analyzed using a Quantibody^®^ Human Cytokine Array 1 kit (RayBiotech, Inc; Cat #: QAH-CYT-1) according to the manufacturer’s instructions.

### Permeability assay

Permeability was assessed by detecting the diffusion of 10 kDa Cascade blue-dextran or 70 kDa Texas red-dextran from the lower blood channel to the upper brain channel. Briefly, ECM medium containing 100 μg/ml dextran was added into the lower blood channel. Two hours later, the media in the upper brain channel were collected. The concentrations of 10 kDa Cascade blue-dextran were determined by fluorescence intensity using a microplate reader system at 400 nm (excitation) and 440 nm (emission). The concentrations of 70 kDa Texas red-dextran were determined by fluorescence intensity using a microplate reader system (excitation: 595 nm; emission: 630 nm). The apparent permeability coefficient (*P_app_*) was calculated according to the equation below ^52,53^:

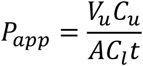

where *V_u_* (ml) and *C_u_* (μg/ml) indicate the volume (100 μl) and concentration of dextran in the upper brain channel, respectively; *A* (cm^2^) indicates the size of transwell membrane (0.336 cm^2^); *C_l_* is the FITC-dextran concentration (100 μg/ml) in the lower blood channel; and *t* (s) is the diffusion time of dextran (7200 s).

### RNA extraction and qRT-PCR

Total RNA was extracted from HBMECs and brain cells separately using TRizol reagent (Invitrogen) as described previously ^54^. cDNA synthesis was performed using PrimeScript RT Master Mix kit (TaKaRa; Cat #: RR036A) and qPCR was performed using TB Green Premix Ex Taq II kit (TaKaRa; Cat #: RR820A). Primers corresponding target genes were showed in Supplementary Table 1. GAPDH was used as reference gene.

### Transmission electron microscopy

After radiation exposure for 4 days, transwells with cells were picked from the fluidic plate and fixed in PBS buffer containing 2.5% glutaraldehyde (Electron Microscopy Sciences) at 4℃ overnight. After washed with PBS for three times and fixed in 1% OsO4 buffer for 2 hours, the samples were dehydrated with graded ethanol solutions, and then embedded in Epon812 resin (SPI). Ultrathin sections (70 nm) were stained with 2% uranyl acetate for 30 minutes and then lead citrate for 10 minutes. Images of the HBMECs were acquired by JEM-1400PLUS electron microscope.

### Measurement of mitochondrial membrane potential

Mitochondrial membrane potential was measured using tetramethylrhodamine methyl ester (TMRM) (Invitrogen). Briefly, cells on transwell were incubated with 20 nM TMRM at 37°C. 20 min, cells were washed two times with PBS and then cultured in corresponding medium. Cells were imaged with using a Leica STELLARIS 5 confocal fluorescent microscope (excitation: 549 nm; emission: 574 nm).

### Western blot

After radiation exposure for 4 days, HBMECs and brain cells were collected and lysed from the transwells, separately. Protein samples were separated on 10% SDS-PAGE and then transferred onto 0.2 μm nitrocellulose membranes (GE Amersham). After being blocked with 5% BSA in TBST buffer containing 0.05% Tween-20, the membranes were probed with the primary antibodies at 4 ℃ overnight. The membranes were then probed with corresponding horseradish peroxidase (HRP)-conjugated secondary antibodies at room temperature for 1 hour at room temperature. Protein bands were detected by Prime Western Blotting Detection Reagent (GE life).

### Drug testing

4 hours before radiation (16 Gy), BBB MPSs were treated with indicated drugs, including 1 μM Hydrocortisone (StemCell; Cat #: 07926), Crisaborole (MCE; Cat #: HY-10978), 1 μM Abrocitinib (MCE; Cat #: HY-107429), 10 μM Coenzyme Q10 (MCE; Cat #: HY-N0111), 2 μM BAI1 (MCE; Cat #: HY-103269) and 20 μM Idebenone (MCE; Cat #: HY-N0303). 4 days after radiation exposure, control and irradiated BBB samples were examined by dextran permeability assay, RNA-seq and immunofluorescence.

### RNA extraction, library preparation and sequencing

Four days after radiation exposure, brain endothelial cells and brain cells (pericytes, astrocytes and microglia) were collected from the lower vascular channel and upper brain channel, respectively. Total RNAs were extracted using TRIzol (Invitrogen) following the methods of Chomczynski et al. ^55^. RNA samples were treated with DNaseI for DNA digestion and then determined by examining A260/A280 using a Nanodrop OneC spectrophotometer (Thermo Fisher Scientific Inc). RNA integrity was confirmed by 1.5% agarose gel electrophoresis. Finally, qualified RNAs were quantified by Qubit 3.0 with a Qubit RNA Broad Range Assay kit (Life Technologies).

Samples with 2 μg total RNA were used for stranded RNA sequencing library preparation using a KC-Digital Stranded mRNA Library Prep Kit for Illumina (Catalog NO. DR08502, Wuhan Seqhealth Co., Ltd. China), following the manufacturer’s instructions. The kit eliminates duplication bias in PCR and sequencing steps by using a unique molecular identifier (UMI) of 8 random bases to label the pre-amplified cDNA molecules. The library products corresponding to 200–500 bps were enriched, quantified, and finally sequenced on a Hiseq X 10 sequencer (Illumina).

### RNA-sequences analysis

Bulk RNA-seq was conducted by Seqhealth Technology Co., LTD (Wuhan, China) using DNBSEQ-T7. Firstly, to discard the low-quality reads and adaptor sequences, raw sequencing data was filtered by Trimmomatic (version 0.36). Subsequently, in-house scripts were further processed using custom scripts to mitigate duplication bias introduced during library preparation and sequencing. In summary, clean reads were initially clustered based on unique molecular identifiers (UMIs), grouping reads with identical UMI sequences. Within each cluster, reads underwent pairwise alignment, and those with over 95% sequence identity were consolidated into new sub-clusters. Multiple sequence alignment was then applied to each sub-cluster to derive a consensus sequence, effectively minimizing errors and biases introduced by PCR amplification and sequencing.

Standard RNA-seq analysis was applied to treat the deduplicated consensus sequences. They were mapped to the reference genome of Homo sapiens from the Ensembl database (ftp://ftp.ensembl.org/pub/release-87/fasta/homo_sapiens/dna/) using STAR aligner v.2.5.3a (Default parameters). The followed data processing was performed using R project (R version 4.4.0). Unique gene hit counts were obtained using the featureCounts function from the Subread package (version 1.5.2) and then RPKM was calculated. Differential expression analysis was then identified using the edgeR package (version 3.12.1). Significant DEGs were defined with p-value <0.05 and the absolute value of log2(fold change) >1 as the threshold. GO and KEGG enrichment analysis of DEGs were implemented using KOBAS software (v.2.1.1) with P value cut-off of 0.05 to determine statistical enrichment. GSEA enrichments were analyzed using GSEA function from the clusterProfiler (version 4.14.0) by the p-value cutoff of 0.05 ^56^. Reactome pathway enrichment analysis was analyzed using the ReactomePA packages (version 1.50.0) ^57^.

### scRNA-sequences analysis

The raw data of single-cell RNA sequences was downloaded from the National Genomics Data Center (NGDC: HRA003477) of China. Human brain tissue samples from patients (*n* = 2) with were obtained from patients with radiation-induced brain injury (RIBI) following cranial radiotherapy for nasopharyngeal carcinoma (NPC). Tissues were categorized as distal or proximal based on their location relative to the lesion core. Additional brain tissues were collected during neurosurgery from patients (*n* = 2) with epilepsy or cerebral hemangiomas, where samples consisted of surgical disposals.

Following data acquisition, FASTQ files were processed using Cell Ranger toolkit (version 8.0.1; 10x Genomics). Raw sequencing reads were aligned to the human reference genome (GRCh38). Default parameters were used for alignment, barcodes assignment and UMI quantification to obtain a filtered matrix containing normalized gene counts versus cells per sample, which were used for downstream analyses.

Data preprocessing was conducted using the Seurat (version 5.1.0) packages. Mitochondrial gene content was quantified (percent.mt), with each dataset filtered to exclude low-quality cells. Specifically, cells with fewer than 300 unique genes or with a mitochondrial gene content exceeding 20% were excluded. Quality control (QC) metrics were visualized pre-filtering to ensure data integrity, using violin plots that summarize feature counts, RNA counts, and mitochondrial gene content for each sample.

Each Seurat object was subjected to a standardized normalization process using the NormalizeData function, followed by identification of the top 2000 variable features. Principal component analysis (PCA) was performed on variable genes, and the top 20 components were selected to capture major sources of variation in gene expression. Cell clustering was conducted by identifying nearest neighbors (FindNeighbors) and applying a clustering algorithm (FindClusters, resolution = 0.5). Uniform Manifold Approximation and Projection (UMAP) was used for low-dimensional representation. To mitigate technical artifacts, DoubletFinder was applied with an 8% doublet formation rate to identify and remove doublets ^58^.Given the multi-sample setup, batch effects were corrected using the Harmony algorithm ^59^. This harmonized dataset was further clustered using Harmony-corrected principal components, allowing a seamless integration of multiple experimental conditions while preserving biological variability. Cell clusters were visualized using UMAP and tSNE, with cells grouped by both sample and inferred cell type. Marker-based annotation of clusters was performed to support downstream biological interpretation.

### Statistical analyses

Data were collected and organized by Microsoft Excel software. GraphPad Prism 8 software was utilized for data statistical analysis. All experiments (except RNA-seq experiment) were repeated at least three times (≥ 3 biological replicates). During the experiment and assessing the outcome, the investigators were blinded to the group allocation. Differences between two groups were analyzed using a Student’s t-test. Multiple group comparisons were performed using a one-way analysis of variance (ANOVA) followed with Bonferroni post hoc test. Data are presented as the mean ± standard error of the mean (SEM). Significance is indicated by asterisks: *: *P* < 0.05, **: *P* < 0.01, ***: *P* < 0.001.

## Data availability

The main data supporting the results in this study are available within the paper and its Supplementary Information. The RNA-seq raw data have been deposited on Sequence Read Archive (SRA) under the accession number GSE280303 and GSE280304. All data generated or analysed during the study are available from the corresponding author on reasonable request.

## Acknowledgements

The authors would like to acknowledge Dr. Wenjie Li (State key Laboratory of Radiation and Protection, Soochow University) for assisting radiation exposure experiments. The authors would like to thank Yingqi Guo in Kunming Biological Diversity Regional Center of Instruments, Kunming Institute of Zoology, CAS, for her help in TEM work. The authors would like to thank Prof. Lin Jin (Shanxi Agricultural University) for kindly providing the antibodies to detect the cGAS-STING pathway, and Prof. Andy Peng Xiang (Sun Yat-Sen University) for kindly providing the hPSC-PC cells. The authors would like to acknowledge Prof. Yamei Tang (Sun Yat-Sen Memorial Hospital) for sharing the scRNA-seq data of patients’ samples. The authors would like to acknowledge Dr. Peng Wang in the Biomedical Platform at Suzhou Institute for Advanced Research of USTC for technical support in bioimaging experiments.

This research was supported by the National Key R&D Program of China (No. 2022YFA1104700), National Nature Science Foundation of China (No. 32101163, 31971373).

## Notes

### Competing Interest Statement

The authors have declared no competing interest.

