## Supplementary Information for "Radiation exposure induced blood-brain barrier injury via mitochondria-mediated sterile inflammation"

**Supplementary Table 1. Primer sequences used for RT-qPCR analysis in this study.**

| Primer | Sequence (5'-3') |
|---|---|
| GAPDH-F | CACCCACTCCTCCACCTTTGAC |
| GAPDH-R | GTCCACCACCCTGTTGCTGTAG |
| IFNE-F | GGAACTGTGTTGGTGCTGCTG |
| IFNE-R | TGGTAGACACTGCTGAATTGACAAG |
| IFNAR1-F | ACTCATTTACACCATTTCGCAAAGC |
| IFNAR1-R | ACCATCCAAAGCCCACATAACAC |
| IFNAR2-F | CCACTCCATTGTACCAACTCACTATAC |
| IFNAR2-R | GCACAGTTCTTAACCACCTTCAAATC |
| STAT1-F | ATGCTGGCACCAGAACGAATGAG |
| STAT1-R | TCACCACAACGGGCAGAGAGG |
| MX1-F | GCATCTCCAGCCACATCCCTTTG |
| MX1-R | TGGTGTCGCTCCGCTCCTTC |
| ISG15-F | TGGACAAATGCGACGAACCTCTG |
| ISG15-R | GCCCGCTCACTTGCTGCTTC |
| IRF7-F | AGAAGAGCCTGGTCCTGGTGAAG |
| IRF7-R | AGGCTGAGGCTGCTGCTATCC |
| IFIT3-F | TACGCCTGGGTCTACTATCACTTGG |
| IFIT3-R | CACTTCAGTTGTGTCCACCCTTCC |
| MT-ND1-F | CCACCTCTAGCCTAGCCGTTTA |
| MT-ND1-R | GGGTCATGATGGCAGGAGTAAT |

**Supplementary Table 2. Primary antibodies used for immunofluorescence in this study.**

| Antibody | Vendor | Catalog # | Dilution |
|---|---|---|---|
| Anti-VE-cadherin | Cell Signaling Technology | 2500S | 1:200 |
| Anti-GFAP | Cell Signaling Technology | 3670S | 1:200 |
| Anti-IBA1 | Proteintech Group | 66827-1-Ig | 1:200 |
| Anti-IBA1 | Proteintech Group | 26177-1-AP | 1:200 |
| Anti-CD31 | HUABIO | M1511-8 | 1:200 |
| Anti-p-STAT1 | Proteintech Group | 28977-1-AP | 1:200 |
| Anti-Ki67 | Abcam | ab243878 | 1:200 |
| Anti-53BP1 | Santa Cruz | sc-515841 | 1:100 |
| Anti-53BP1 | Abcam | ab175933 | 1:200 |
| Anti-dsDNA | Abcam | ab27156 | 1:200 |
| Anti-cGAS | Proteintech Group | 26416-1-AP | 1:200 |
| Anti-TOM20 | Proteintech Group | 11802-1-AP | 1:200 |

**Supplementary Table 3. Primary antibodies used for Western blot in this study.**

| Antibody | Vendor | Catalog # | Dilution |
|---|---|---|---|
| Anti-p-TBK1 | Cell Signaling Technology | 5483 | 1:2000 |
| Anti-TBK1 | Cell Signaling Technology | 3504 | 1:2000 |
| Anti-cGAS | Proteintech Group | 26416-1-AP | 1:2000 |
| Anti-STING | Proteintech Group | 19851-1-AP | 1:2000 |
| Anti-GAPDH | Proteintech Group | 60004-1-Ig | 1:2000 |

**Supplementary Table 4. Chemicals used in this study.**

| Antibody | Vendor | Catalog # | Final concentration |
|---|---|---|---|
| Hydrocortisone | Stemcell | 07925 | 1 μM |
| Crisaborole | MedChemExpress | HY-10978 | 10 μM |
| Abrocitinib | MedChemExpress | HY-107429 | 1 μM |
| SN-011 | MedChemExpress | HY-145010 | 1 μM |
| Coenzyme Q10 | MedChemExpress | HY-N0111 | 10 μM |
| BAI1 | MedChemExpress | HY-103269 | 2 μM |
| Idebenone | MedChemExpress | HY-N0303R | 20 μM |

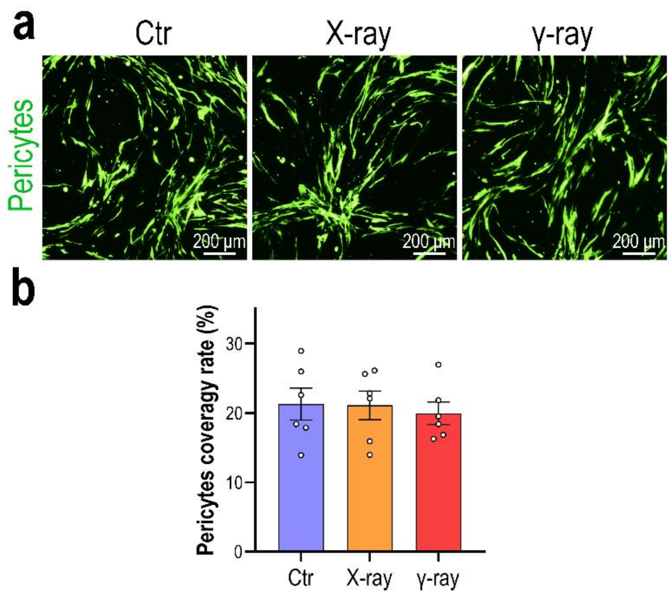


**Supplementary Figure 1. Fluorescent image showing pericytes on BBB MPS following radiation exposure. a**, Fluorescent image showing pericytes labelled with GFP on BBB MPS, 4 days following radiation exposure. **b**, Quantification of pericytes coverage rate based on **a** (*n* = 3). Two images were analyzed for each sample. Data are presented as the mean ± SEM and were analyzed using a one-way analysis of variance (ANOVA) followed by the Bonferroni post hoc test.

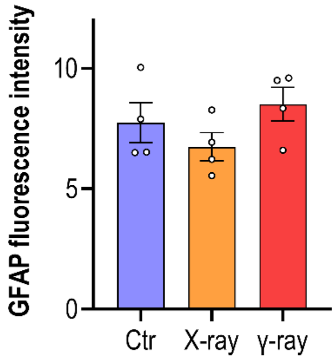


**Supplementary Figure 2. Quantification of GFAP fluorescence intensity for astrocytes on BBB MPS following radiation exposure.** Quantification of GFAP fluorescent intensity based on **Fig. 2f** (*n* = 4). Data are presented as the mean ± SEM and were analyzed using a one-way analysis of variance (ANOVA) followed by the Bonferroni post hoc test.

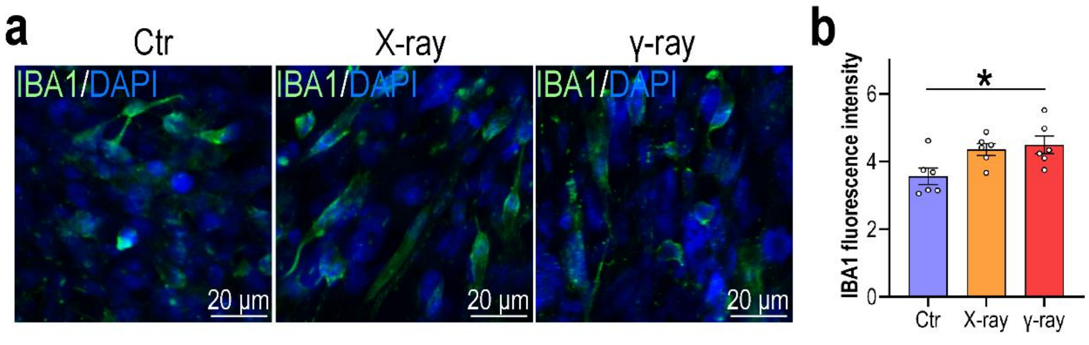

**Supplementary Figure 3. Immunofluorescent image showing microglia on BBB MPS following radiation exposure on BBB MPS. a**, Confocal micrographs showing microglia immunostained for IBA1 on the BBB MPS, 4 days after radiation exposure ($n$ = 3). **b**, Quantification of IBA1 fluorescent intensity based on **a**. Two images were analyzed for each sample. Data are presented as the mean ± SEM and were analyzed using a one-way analysis of variance (ANOVA) followed by the Bonferroni post hoc test (*: $P < 0.05$).

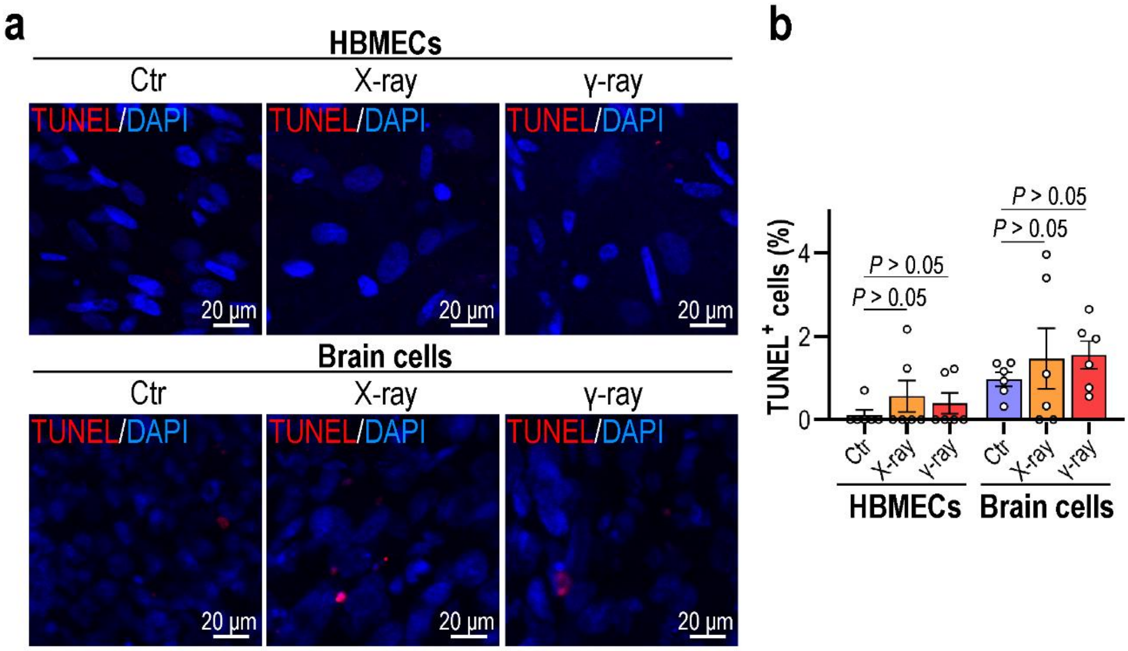


**Supplementary Figure 4. Apoptosis analysis for BBB MPS following radiation exposure. a**, Confocal micrographs showing cells stained by TUNEL kit on the BBB MPS, 4 days after radiation exposure ($n$ = 3). **b**, Quantification of TUNEL+ cells based on **a**. Two images were analyzed for each sample. Data are presented as the mean ± SEM and were analyzed using a one-way analysis of variance (ANOVA) followed by the Bonferroni post hoc test.

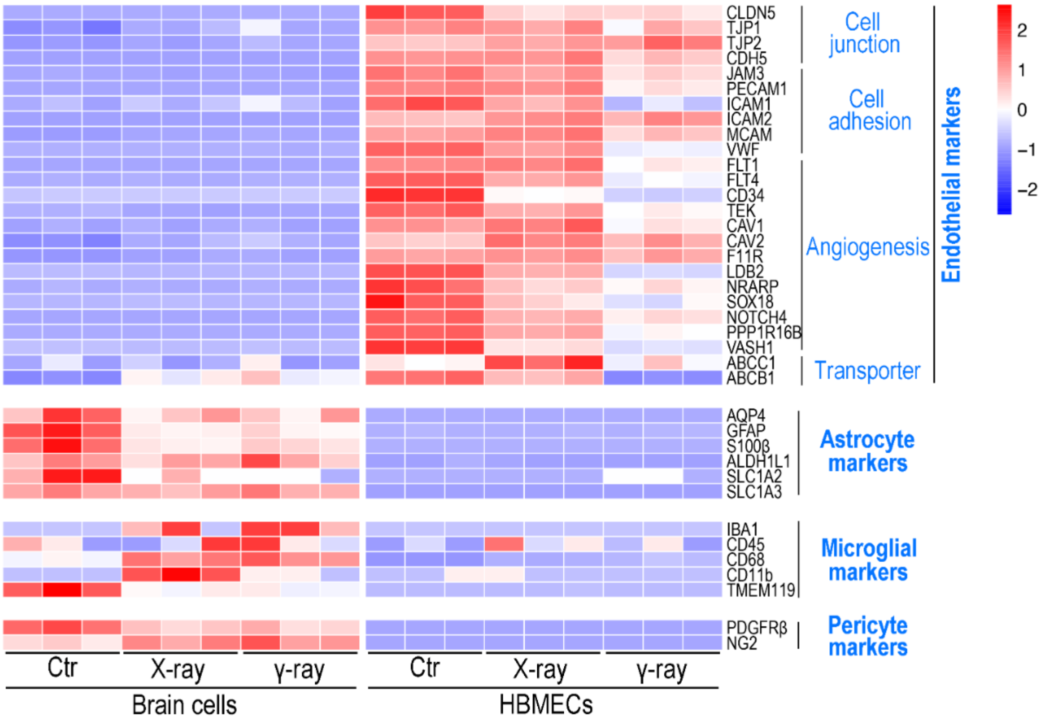


**Supplementary Figure 5. Transcriptomic analysis showing expression levels of cell-specific genes for BBB MPS following radiation exposure.** Heatmap showing the expression of cell-specific genes in brain endothelial cells, astrocytes, microglia and pericytes, 4 days after radiation exposure ($n$ = 3). Genes differentially expressed with fold changes of > 2.0 and $P$ < 0.05 are defined as differentially expressed genes (DEGs). $P$ values were calculated using a two-sided, unpaired Student's t-test with equal variance assumed.

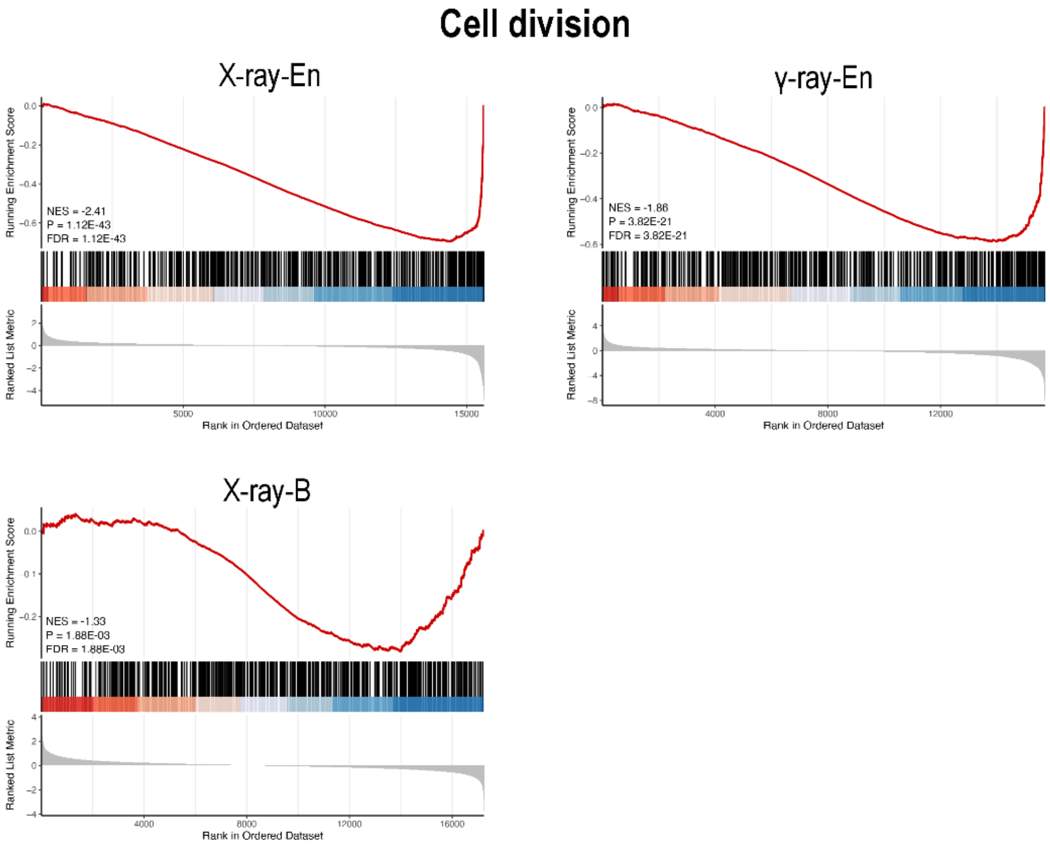


**Supplementary Figure 6. GSEA analysis of cell division process for brain endothelial cells and brain cells on the BBB MPS following radiation exposure. a**, GSEA analysis reveals correlation between radiation exposure and genes involved in cell division in brain endothelial cells. **b**, GSEA analysis reveals correlation between radiation exposure and genes involved in cell division in brain cells. **a-b**, NES, normalized enrichment score. FDR, false discovery rate. Gene sets were considered significant when $P < 0.05$ and FDR < 0.25.

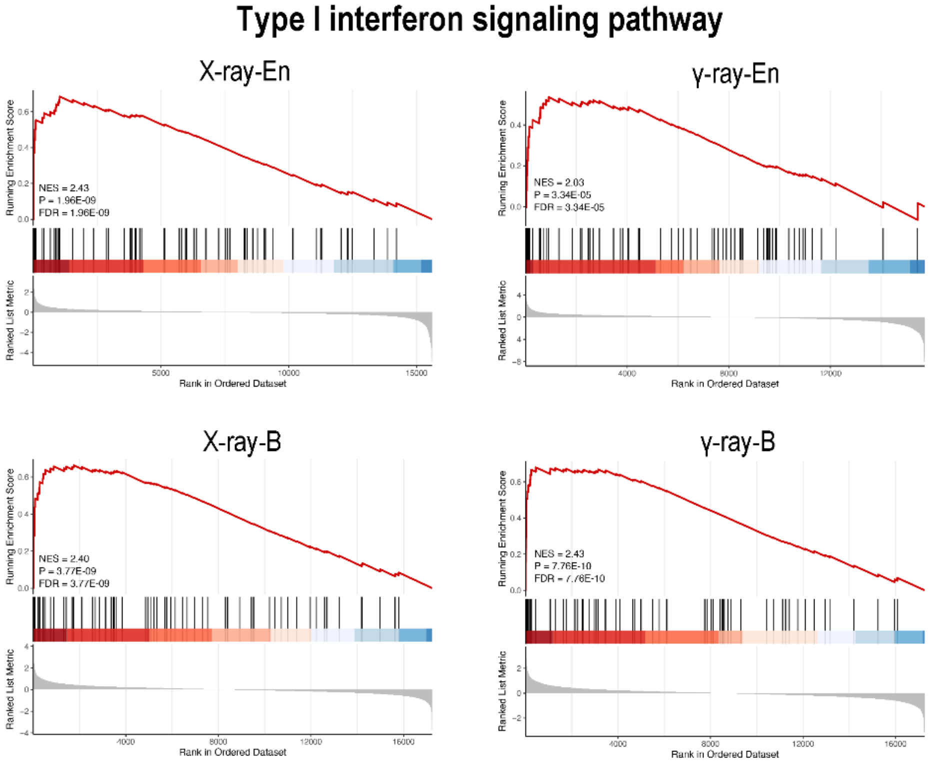


**Supplementary Figure 7. GSEA analysis of type I interferon signaling pathway for brain endothelial cells and brain cells on the BBB MPS following radiation exposure. a**, GSEA analysis reveals correlation between radiation exposure and genes involved in type I interferon signaling pathway in brain endothelial cells. **b**, GSEA analysis reveals correlation between radiation exposure and genes involved in type I interferon signaling pathway in brain cells. **a-b**, NES, normalized enrichment score. FDR, false discovery rate. Gene sets were considered significant when $P < 0.05$ and FDR < 0.25.

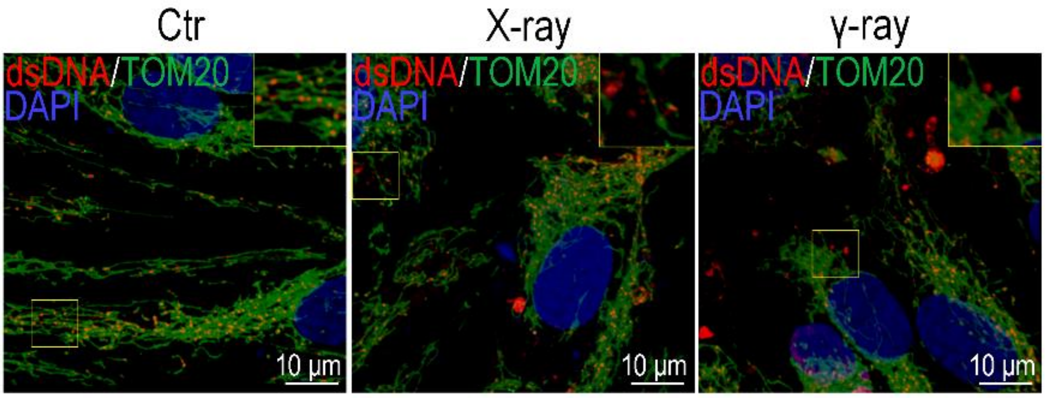


**Supplementary Figure 8. 3D micrographs showing sub-cellular distribution of dsDNA and TOM20 in brain endothelial cells following radiation exposure.** 3D micrographs showing brain endothelial cells immunostained for TOM20 (green) and dsDNA (red) in brain endothelial cells, 4 days after radiation exposure.

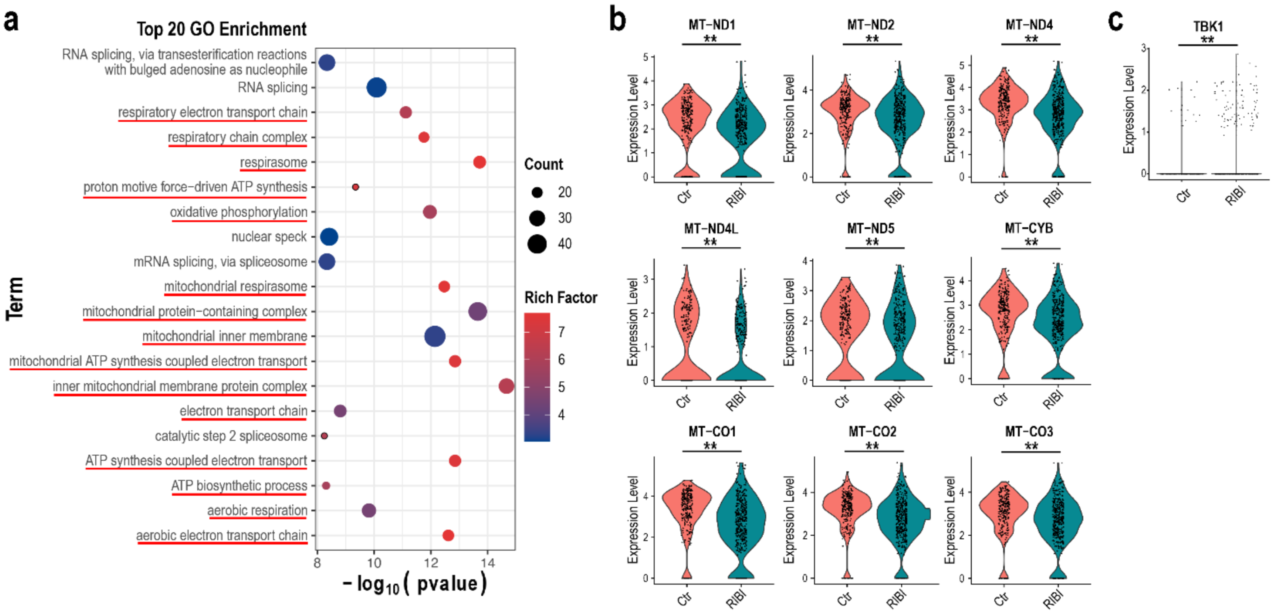


**Supplementary Figure 9. scRNA-seq analysis for brain endothelial cells from RIBI patients. a**, GO enrichment analysis of down-regulated DEGs in brain endothelial cells from RIBI patients' samples. **b**, Violin plots showing levels of mtDNA-encoded genes in brain endothelial cells from RIBI patients' samples. **c**, Violin plots showing level of TBK1 in brain endothelial cells from RIBI patients' samples. The scRNA-seq data was obtained from a study of Tang's group (Shi et al., 2023).

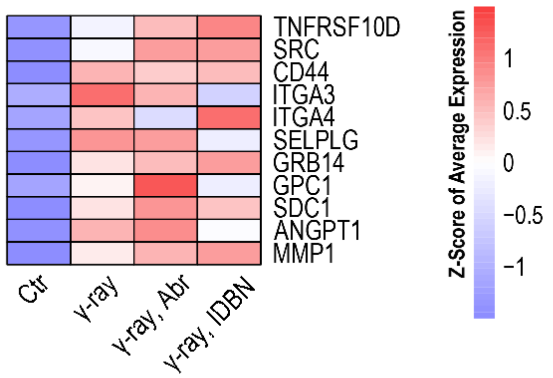


**Supplementary Figure 10. Heatmap showing expression levels of genes related to leukocyte migration in irradiated brain endothelial cells following abrocitinib or idebenone treatment.** Genes differentially expressed with fold changes of > 2.0 and $P < 0.05$ are defined as differentially expressed genes (DEGs). $P$ values were calculated using a two-sided, unpaired Student's t-test with equal variance assumed ($n = 3$).

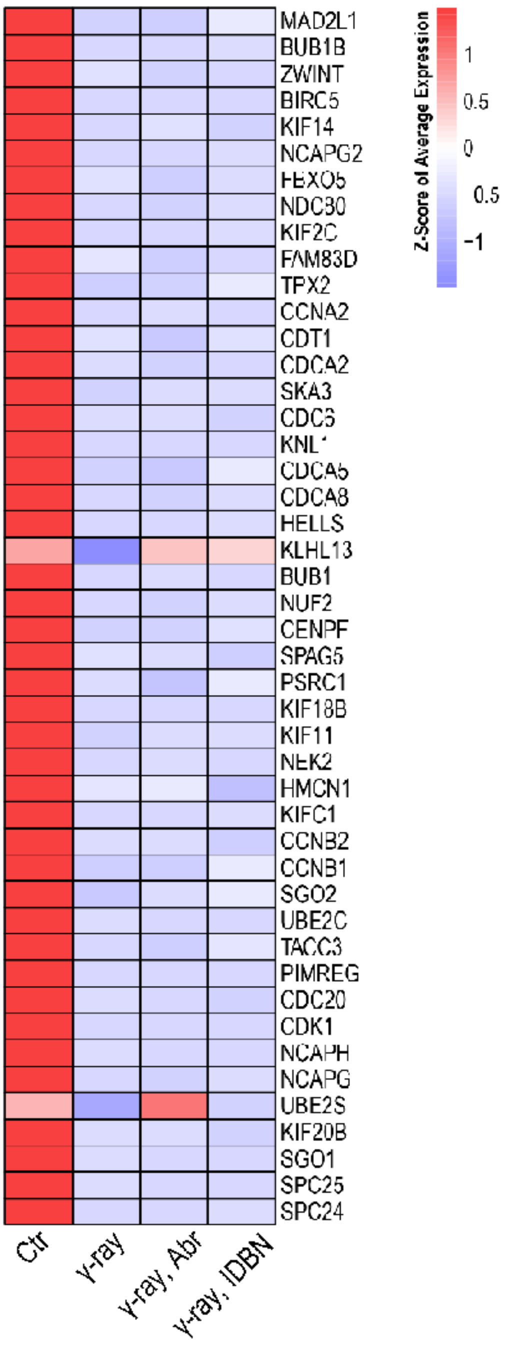


**Supplementary Figure 11. Heatmap showing expression levels of genes related to cell division in irradiated brain endothelial cells following abrocitinib or idebenone treatment.** Genes differentially expressed with fold changes of > 2.0 and *P* < 0.05 are defined as differentially expressed genes (DEGs). *P* values were calculated using a two-sided, unpaired Student's t-test with equal variance assumed (*n* = 3).

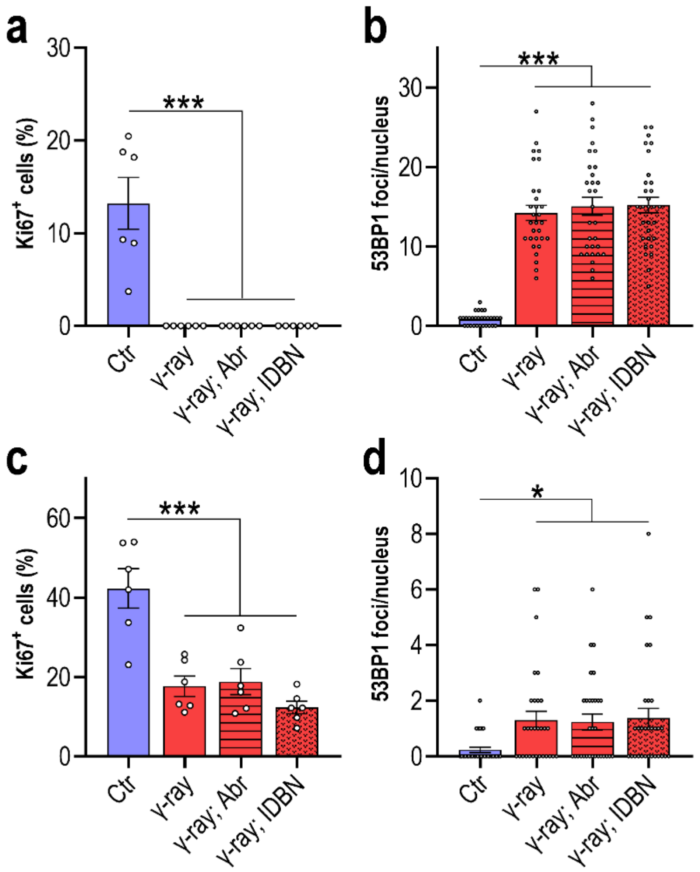


**Supplementary Figure 12. Quantification of Ki67+ cells and 53BP1 foci for cells of irradiated BBB MPS following abrocitinib or idebenone treatment. a**, Quantification of Ki67+ cells for brain endothelial cells on irradiated BBB MPS following abrocitinib or idebenone treatment based on **Fig. 6j**. **b**, Quantification of 53BP1 foci per cell for brain endothelial cells on irradiated BBB MPS following abrocitinib or idebenone treatment based on **Fig. 6j**. **c**, Quantification of Ki67+ cells for brain cells on irradiated BBB MPS following abrocitinib or idebenone treatment based on **Fig. 6l**. **d**, Quantification of 53BP1 foci per cell for brain cells on irradiated BBB MPS following abrocitinib or idebenone treatment based on **Fig. 6l**. Data are presented as the mean ± SEM and were analyzed using a one-way analysis of variance (ANOVA) followed by the Bonferroni post hoc test (*: $P < 0.05$; ***: $P < 0.001$).
